# Identifying Conserved Genomic Elements and Designing Universal Probe Sets To Enrich Them

**DOI:** 10.1101/077172

**Authors:** Brant C. Faircloth

## Abstract

Targeted enrichment of conserved genomic regions is a popular method for collecting large amounts of sequence data from non-model taxa for phylogenetic, phylogeographic, and population genetic studies. Yet, few open-source workflows are available to identify conserved genomic elements shared among divergent taxa and to design enrichment baits targeting these regions. These shortcomings limit the application of targeted enrichment methods to many organismal groups. Here, I describe a universal workflow for identifying conserved genomic regions in available genomic data and for designing targeted enrichment baits to collect data from these conserved regions. I demonstrate how this computational approach can be applied to diverse organismal groups by identifying sets of conserved loci and designing enrichment baits targeting thousands of these loci in the understudied arthropod groups Arachnida, Coleoptera, Diptera, Hemiptera, or Lepidoptera. I then use *in silico* analyses to demonstrate that these conserved loci reconstruct the accepted relationships among genome sequences from the focal arthropod orders, and we perform *in vitro* validation of the Arachnid probe set as part of a separate manuscript (Starrett *et al.* Submitted). All of the documentation, design steps, software code, and probe sets developed here are available under an open-source license for restriction-free testing and use by any research group, and although the examples in this manuscript focus on understudied and exceptionally diverse arthropod groups, the software workflow is applicable to all organismal groups having some form of pre-existing genomic information.

---

Collecting sequence data from non-model taxa has undergone a revolution during the previous 10 years, driven by advancements in sequencing technologies (Bentley *et al.* 2008) and molecular methods (Hardenbol *et al.* 2003; Baird *et al.* 2008; Gnirke *et al.* 2009). Ecologists and evolutionary biologists have typically focused on a narrower subset of these approaches, collectively known as “reduced-representation” methods, which include varieties of restriction-enzyme-based (Baird *et al.* 2008; Elshire *et al.* 2011; Peterson *et al.* 2012), transcriptomic (Dunn *et al.* 2008; Smith *et al.* 2011; Misof *et al.* 2014), and targeted enrichment (Faircloth *et al.* 2012; Bi *et al.* 2012; Peñalba *et al.* 2014; Ali *et al.* 2015; Hugall *et al.* 2016; Hoffberg *et al.* 2016; Suchan *et al.* 2016) approaches. These methods allow the collection of large numbers of loci from large numbers of organisms and are less expensive and potentially less complicated than whole-genome sequencing or genome-resequencing approaches, particularly when collecting data from tens or hundreds of individuals.

One popular reduced representation approach is the targeted enrichment (Gnirke *et al.* 2009) of conserved or ultraconserved genomic elements (sensu Faircloth *et al.* 2012). In this approach, researchers identify genomic regions of high conservation shared among divergent lineages, design synthetic oligonucleotide “baits” that are complementary to these regions, hybridize genomic libraries to these oligonucleotide baits, “fish” out the hybridized bait+library structure, remove the bait sequence, and sequence the remaining pool of enriched, targeted DNA. Although the baits target and enrich conserved regions of the genome, the library preparation, enrichment, sequencing, and assembly procedures ensure that the approach also captures variable flanking sequence that sits to each side of each conserved region (Faircloth *et al.* 2012; Smith *et al.* 2014).

The power of this approach is that a single tube of synthetic oligonucleotide baits can be used by multiple studies to collect data across very broad taxonomic scales - for example amniotes (Crawford *et al.* 2012; McCormack *et al.* 2013; Hosner *et al.* 2015; Streicher & Wiens 2016) or fishes (Faircloth *et al.* 2013; McGee *et al.* 2016) or bees, ants, and wasps (Faircloth *et al.* 2015; Blaimer *et al.* 2015). Additionally, the sequence reads from these enriched, conserved loci can be analyzed in different ways to address questions at different scales, from deep-time phylogenetic studies (Faircloth *et al.* 2013) to shallower level phylogeographic studies (Smith *et al.* 2014) to population-level studies (Harvey *et al.* 2016; Manthey *et al.* 2016). Finally, because the targeted enrichment approach is DNA-based, it can be applied to degraded and low-quantity samples, such as those in many specimen or tissue collections (Bi *et al.* 2013; McCormack *et al.* 2015; Lim & Braun 2016; Blaimer *et al.* 2016).

Although targeted enrichment of conserved elements offers many benefits (Harvey *et al.* 2016), there are few explicit, easy-to-use workflows for identifying conserved loci shared among organismal genomes or for designing sequence capture probes targeting these regions (cf. Mayer *et al.* 2016; Johnson *et al.* 2016). Here, I describe the workflow I have recently developed to accomplish these tasks, and I demonstrate its utility by: (1) identifying large suites of conserved elements shared within five diverse and understudied arthropod orders (Arachnida, Coleoptera, Diptera, Hemiptera, Lepidoptera), and (2) designing five sets of capture baits targeting conserved regions shared among members of each taxonomic order. This updated workflow differs from previous approaches (Faircloth *et al.* 2012; McCormack *et al.* 2012) by aligning small, random pieces of DNA from several genomes to a focal reference genome using a permissive read aligner and then using overlapping coordinates shared among multiple taxa to identify regions of shared conservation. This technique greatly increases the number of conserved regions detected relative to synteny based, genome-genome alignment procedures (e.g., (Harris 2007)) used in earlier manuscripts (Faircloth *et al.* 2012; McCormack *et al.* 2012). I then use *in silico* target enrichment experiments to show that these bait sets collect conserved loci that can be used to reconstruct the known phylogenetic relationships within their respective class/orders. We empirically test one of these probe sets by enriching conserved loci from a diverse group of Arachnids as part of a separate manuscript (Starrett *et al.* Submitted).

I make all documentation and computer code for this workflow available under an open-source license, allowing researchers to generalize the approach to other organisms having some genomic data. I also make all of the bait sets for Arachnida, Coleoptera, Diptera, Hemiptera, and Lepidoptera available under a public domain license (CC-0), facilitating restriction-free commercial synthesis, testing, use, and improvement of these probe sets by other research groups interested in phylogenetic, phylogeographic, and population-level analyses of arthropods.

## Methods

### Study Group

The arthropod groups Arachnida, Coleoptera, Diptera, Hemiptera, and Lepidoptera are among the most diverse invertebrate classes/orders, encompassing more than 900,000 species (Harvey 2002; Zhang 2011). Yet, our understanding of the evolutionary factors responsible for generating the extreme diversity within each of these groups is poor. Large projects, like the i5k (i5K Consortium 2013) are transforming our knowledge of major relationships among arthropod lineages (Misof *et al.* 2014). However, the extreme diversity of many arthropod groups makes the large-scale collection of transcriptome data across clades difficult, suggesting that less expensive, genome reduction techniques that work with DNA (versus RNA) could be useful for understanding finer-grained evolutionary relationships among hundreds or thousands of arthropod species within major taxonomic groups. Targeted enrichment of conserved DNA regions shared among these species offers one approach for beginning to fill these gaps, particularly because the technique is useful with older, degraded DNA, similar to that collected from arthropod (Faircloth *et al.* 2015; Blaimer *et al.* 2016) and other museum specimens (Bi *et al.* 2013; McCormack *et al.* 2015).

### General Workflow

Although some implementation details differ for the groups described below in terms of the specific data used, the general workflow (Figure 1) for identifying conserved loci and designing capture baits to target them begins with the selection of an appropriate “base” genome that is within or related to the focus group and against which data from other exemplar taxa sampled within the focus group will be aligned. The base genome sequence can be an ingroup or outgroup taxon, and it is reasonable to select the best-assembled and annotated genome that is closely related to or nested within the focus group rather than focusing intently on ingroup or outgroup status. This choice facilitates downstream analysis or selection of conserved loci based on desirable properties derived from annotation or positional information (exon, intron, intergenic, unlinked, etc.), although how, exactly, to identify the best-assembled genome is a matter of debate (Earl *et al.* 2011; Bradnam *et al.* 2013). I generally focus on selecting assemblies as the “base” when they contain relatively large and complete scaffolds and have reasonable annotation (ideally evidence-based, although gene predictions are also useful).

**Fig 1:**
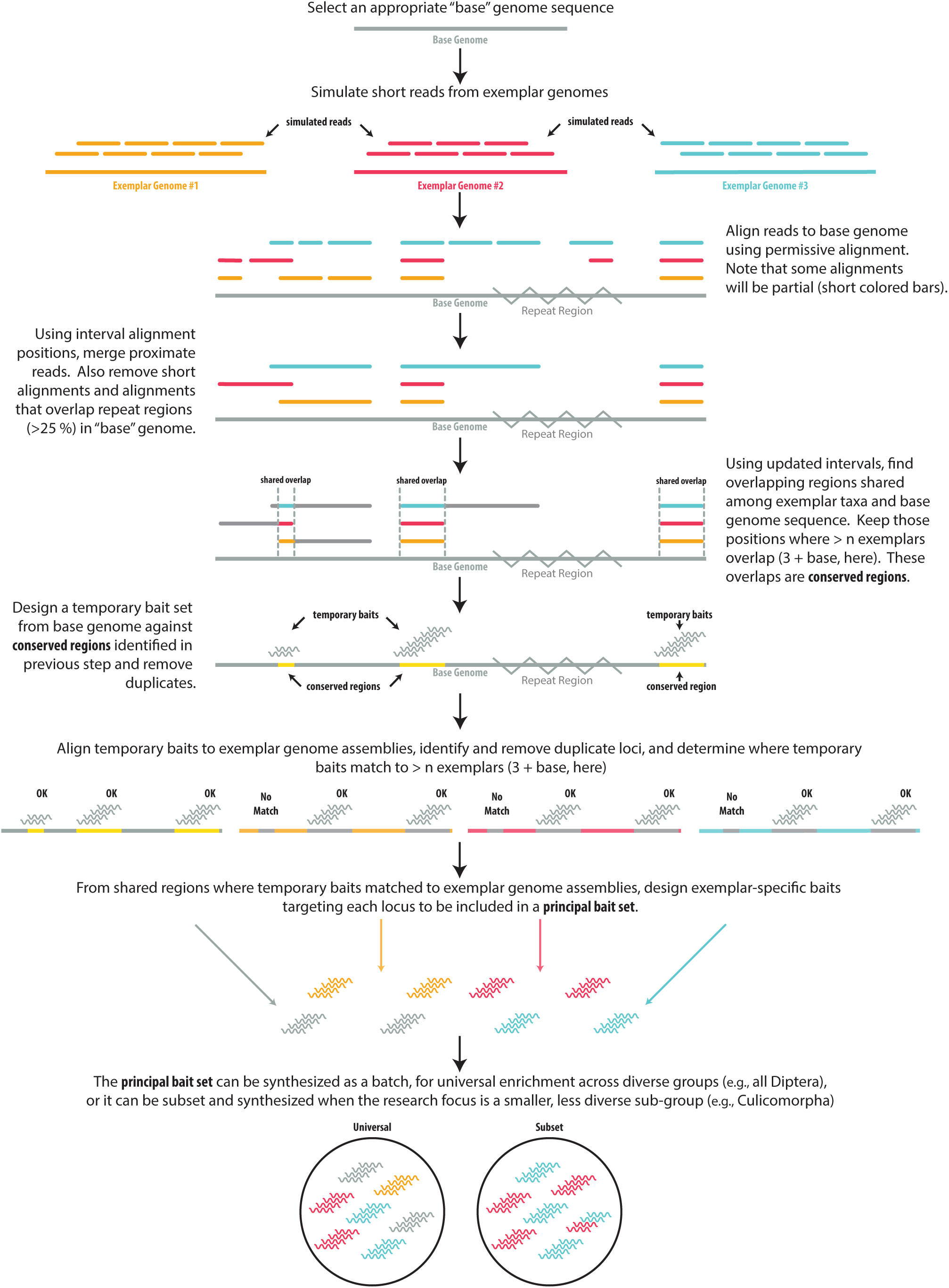
Illustration of the steps involved in the conserved element identification and probe design workflow.

Following the selection of a base genome, the workflow proceeds by generating short reads from organisms that serve as exemplars of the focus group’s diversity using either: (1) low-coverage (4-6x), massively parallel sequencing reads and/or (2) short reads simulated from other genome sequences that exist for the focus group. The next step in the workflow is to align all sets of exemplar reads to the base genome using a permissive raw-read aligner such as *stampy*(Lunter & Goodson 2011), produce a BAM (Li *et al.* 2009) file, and use *samtools* (Li *et al.* 2009) to reduce the size of the BAM file by selecting only the reads from each BAM file that align to the base genome.

The workflow proceeds by converting successfully aligned reads to interval (BED) format using *bedtools* (Quinlan & Hall 2010), which allows fast and easy manipulation of alignment data. Using the resulting BED files, the next step in the workflow is to sort the alignment coordinates; merge together alignment positions in each file that are close (<100 bp) to one another; and remove alignments that are short (<80 bp), overlap masked loci (>25% of length) and/or contain ambiguous (N or X) bases. The workflow then proceeds by processing the filtered BEDs to create a relational database of overlapping alignment positions shared between the base genome sequence and each of the exemplar taxa. Because reads from each exemplar taxon were aligned to the same base genome and because masked loci have been removed, overlapping alignment positions shared among taxa represent loci that are putatively conserved between genomes. Users can query this database to generate a BED file of the genomic locations of each conserved locus in the base genome sequence that are also shared by a subset of some or all of the exemplar taxa.

The final stages of the workflow focus on designing oligonucleotide baits to target the conserved loci identified in the steps described above. The first step of this process is to extract the conserved loci from the base genome as FASTA-formatted records, and design temporary oligonucleotide bait sequences targeting these loci. The workflow then uses LASTZ (Harris 2007) to align these temporary baits designed from the base genome to genomic data from a set of exemplar taxa (which can be the same, a subset, or a superset of the organisms used for locus identification) and builds a relational database of loci detected in each of the exemplar taxa.

From this relational database, users can determine which base genome probe sequences “hit” in which exemplar taxa, and they can select to output those loci consistently detected in a majority of exemplar taxa. Users input this list of loci to a program that designs bait sequences from all exemplar taxa where each conserved locus was consistently detected. The final stage of the workflow is to screen and remove probe sequences that appear to target duplicate loci within and between all exemplar taxa. The final output of the workflow is a file containing bait sequences for each conserved locus that were selected from each exemplar genome, such that Locus 1 may have baits designed from Taxon A, Taxon B, and Taxon C. This design approach increases the likelihood that baits will capture the targeted locus when combined with DNA libraries prepared from organisms having genome sequences divergent from the exemplar taxa. I called this final bait design file the “principal” FASTA file of bait sequences or the “principal bait set”.

### Arachnida

Because few arachnids have genomic data available, I collected sequence data from a diverse group of arachnids using low-coverage genome-sequencing. I extracted DNA from legs or legs+cephalothorax of samples using Qiagen DNeasy kits, adding 2 µL RNase (1 mg/mL) to each extraction. I visualized DNA extracts on 1.5% (w/v) agarose, and I sheared the resulting DNA to 400-500 bp using a Bioruptor (Diagenode, Inc.). After shearing, I prepared sequencing libraries from 100-500 ng sheared DNA using a commercial library preparation kit (Kapa Biosystems, Inc.) with a set of custom sequence tags to identify each library (Faircloth & Glenn 2012). I amplified 15 µL of each library in a reaction mix of 25 µL Kapa HiFi HS Master Mix (Kapa Biosystems, Inc.), 5 µL Illumina primer mix (5 µM each), 15 µL of adapter-ligated DNA, and 5 µL of ddH2O using a thermal profile of 98 °C for 45 seconds followed by 14 cycles of 98 °C for 15 seconds, 60 °C for 30 seconds, 72 °C for 60 seconds followed by a 72 °C extension for 5 minutes. After amplification, I cleaned libraries 1:1 with a SPRI-substitute (Rohland & Reich 2012), and I checked the quality of resulting libraries by visualizing 1 µL of each (5 ng/µL) on a BioAnalyzer (Agilent, Inc.). Because I observed adapter-dimer peaks following initial quality control, I cleaned libraries 1-2 additional times using 1:1 SPRI-substitute. After validating the removal of dimer peaks from the sequencing libraries, I qPCR quantified 5 ng/µL aliquots of each library using a commercial kit (Kapa Biosystems, Inc.), and I combined libraries at equimolar ratios to make a 10 µM pool that I sequenced using PE150 reads on an Illumina HiSeq 2000 (UCSC Genome Technology Center). Once I received sequence data from the sequencing center, I trimmed sequencing reads for adapter contamination and low quality bases using an automated wrapper around trimmomatic (Faircloth 2013; Bolger *et al.* 2014), and I merged read pairs into a single file. To add a lineage representing ticks to the sequence data generated from other arachnids, I used art (Huang *et al.* 2012) to simulate error-free, paired-end reads of 100 bp from the *Ixodes scapularis* genome assembly (GCA_000208615.1).

Following the steps outlined in Supplemental File 1, I aligned reads for all organisms in Supplemental Table 1 to the *Limulus polyphemus* genome assembly (GCA_000517525.1; hereafter limPol1) using *stampy* (Lunter & Goodson 2011) with a substitution rate of 0.10, and I streamed the resulting SAM data to a BAM file using *samtools* (Li *et al.* 2009). After alignment, I reduced the BAM file to contain only those reads mapping to the limPol1 genome, I converted each BAM file to a BED file, and I screened the resulting interval data to remove those intervals in each BED file that overlapped masked, short (<80 bp), or ambiguous segments of the limPol1 genome. The intervals that were not filtered represent conserved sequence regions shared between the base genome (limPol1) and each of the exemplar taxa. I used the phyluce_probes_get_multi_merge_table program to determine which of these conserved intervals were shared among some/all of the exemplar taxa, and I output a list of those intervals shared by limPol1 and six arachnid exemplars. I buffered these intervals shared by limPol1 and arachnids to 160 bp, and I extracted FASTA sequence from the limPol1 genome corresponding to the buffered intervals (phyluce_probes_get_genome_sequences_from_bed). Then, I designed a temporary set of sequence capture baits by tiling bait sequences over each interval (phyluce_probe_get_tiled_probes) at 3x density (probes overlapped by 40 bp). This produced a set of temporary enrichment baits designed from limPol1, and I screened this set of temporary baits to remove baits sequences that were ≥50% identical over >50% of their length.

To design a more diverse bait set that included baits from a larger selection of arachnids, I downloaded several arachnid genome assemblies (Supplemental Table 1) and also included a new genome assembly from a tick (NCBI Submission *In Progress*). Then, I aligned baits from the temporary bait set to each genome using a wrapper (phyluce_probe_run_multiple_lastzs_sqlite) around *lastz* (Harris 2007) with liberal alignment parameters (≥50% sequence identity required to map). Using the alignment data, I removed loci that were hit by baits targeting different conserved regions or multiple loci that were hit by the same bait (phyluce_slice_sequence_from_genomes), and I buffered remaining, non-duplicate loci to 180 bp. I used a separate program (phyluce_probes_get_multi_fasta_table) to determine which loci I detected across the arachnid genome assemblies, and I created a list of those loci detected in 6 of the 10 arachnid genome assemblies. I then designed a bait set targeting these loci by tiling probes across each locus in each of the 10 arachnid genomes where I detected the locus, and I screened the resulting bait set to remove putative duplicates. I called this the principal arachnid bait set.

To check the sanity of the data returned from the principal arachnid bait set, I performed an *in silico* targeted enrichment experiment. First, I aligned the baits to 10 arachnid genomes (Supplemental Table 1) using a program (phyluce_probe_slice_sequence_from_genomes) from the PHYLUCE package. After identifying conserved loci that aligned to baits in the principal bait set, I buffered the match locations by ± 500 base pairs, and I extracted FASTA data from the buffered intervals. Then, I input the FASTA-formatted contigs from the previous step to the standard PHYLUCE workflow for phylogenomic analyses (Faircloth 2015). Briefly, I performed additional orthology and duplicate screening steps (phyluce_assembly_match_contigs_to_probes; --min_coverage 80, --min_identity 80), exported non-duplicate conserved loci to FASTA format, aligned the FASTA data using *mafft* (Katoh & Standley 2013), and trimmed the resulting alignments using *gblocks* (Castresana 2000; Talavera & Castresana 2007). I created a dataset in which all alignments contained at least 7 of the 10 taxa (70% complete matrix), and I concatenated the resulting alignment data into a supermatrix. I used *RAxML* v8.0.19 (Stamatakis 2014) to: (1) perform a maximum likelihood (ML) search for the tree best-fitting the data using the GTRGAMMA site rate substitution model, (2) perform non-parametric bootstrapping of the data, and (3) reconcile the “best” ML tree with the bootstrap support values.

To further assess the performance of the principal arachnid bait set, we performed extensive *in vitro* enrichments of the identified loci as part of a separate manuscript (Starrett *et al.* Submitted).

### Coleoptera

To design probes targeting conserved loci in Coleoptera (specific steps are outlined in Supplemental File 2), I downloaded available genomes for several coleopteran lineages (Supplemental Table 2), and I used *art* (Huang *et al.* 2012) to simulate error-free, paired-end reads of 100 bp at 2x coverage from each genome sequence. I merged paired reads for each taxon into a single file, and I aligned the merged, simulated reads to the genome sequence of *Tribolium castaneum* (GCA_000002335.2; triCas1 hereafter) using *stampy* (Lunter & Goodson 2011) with a substitution rate of 0.05 and streaming the resulting SAM alignment data to BAM format using *samtools* (Li *et al.* 2009). Subsequent processing steps were similar to the workflow for Arachnida. In brief, I remove unaligned reads from the BAM file, converted the BAM file to BED format, and I screened the resulting interval data to remove those intervals in each BED file that overlapped masked, short (<80 bp), or ambiguous segments of the triCas1 genome. I subsequently created a table of conserved regions shared between the base genome (triCas1) and each of the exemplar taxa, and I queried this table to output a list of intervals shared by all of the exemplar taxa and the base taxon. I selected this stricter threshold (relative to those used for other arthropod groups) because of the extreme diversity of the beetle clade and because conserved loci shared among all of the exemplar beetle taxa were more likely to be present in all beetle lineages. I output the list of these loci, designed a temporary bait set using FASTA data from the triCas1 genome, and re-aligned the temporary baits to the genomes of each exemplar taxon (Supplemental Table 2), as well as one species representing a strepsipteran outgroup to beetles (*Mengenilla moldrzyki*, GCA_000281935.1). I included this species to add additional diversity to the bait set and better represent earlier diverging clades in the beetle tree relative to the clades represented by the other beetle genomes I used for bait design. I extracted sequence in FASTA format for each conserved locus from each exemplar taxon assembly, and I designed a hybrid set of bait sequences targeting each of these loci from the genomes of the exemplar taxa. I filtered putative duplicate baits/loci from this data set, and I called the resulting file the principal coleopteran bait set.

I performed an *in silico* sanity check of the bait set using an approach identical to that described above. I aligned the principal bait set to the genomes of the taxa that I used to design the principal bait set, sliced FASTA sequences from each genome that flanked the conserved locus location by ± 400 bp, performed additional orthology and duplicate screening steps (-- min_coverage 67, --min_identity 80), used *mafft* to align FASTA slices for each locus across all taxa, trimmed resulting alignments using *gblocks*, created a dataset in which all alignments contained at least 5 of the 7 taxa (70% complete matrix), and concatenated these into a PHYLIP supermatrix which I analyzed using best ML (GTRGAMMA) and bootstrap searches in *RAxML* v8.0.19 (Stamatakis 2014). I reconciled the best ML tree with the bootstrap replicates using *RAxML* v8.0.19.

### Diptera

The probe design process for dipterans (Supplemental File 3) followed the same workflow I used to design the principal coleopteran bait set. I downloaded available genomes for several dipteran lineages (Supplemental Table 3), and I simulated error-free, paired-end, 100 bp reads for conserved locus identification from several genomes at 2x coverage using *art* (Huang *et al.*2012). After merging read pairs into a single file, I aligned the simulated reads to the genome sequence of *Aedes aegypti* (aedAeg1 hereafter) using *stampy* (Lunter & Goodson 2011) with a substitution rate of 0.05 and streaming conversion of the output SAM file to BAM format. After removing unaligned reads from the BAM file, I converted the BAM to a BED file, sorted the BED file, merged alignment intervals that were <100 bp from one another, and screened the resulting interval data to remove intervals that overlapped masked, short (<80 bp), or ambiguous segments of the aedAeg1 genome. I created a table of conserved regions shared between the base genome (aedAeg1) and each of the exemplar taxa, and I queried this table to output a list of intervals shared by all of the exemplar taxa and the base taxon. I output the list of these loci, designed a temporary bait set from the aedAeg1 genome, and re-aligned the temporary baits to the genomes of exemplar taxa representing a diverse group of dipteran species (Supplemental Table 3). I extracted sequence in FASTA format for each conserved locus from each exemplar taxon assembly, designed a set of hybrid bait sequences targeting each of these loci from the genomes of each exemplar taxa, and filtered putative duplicate baits/loci from this set. I called the resulting file the principal dipteran bait set.

I performed an *in silico* check of the bait set by reconstructing the relationships between members of two dipteran clades (Culicomorpha and Drosophilidae), where relationships have previously been resolved with reasonable support (Drosophila 12 Genomes Consortium *et al.*2007; van der Linde *et al.* 2010; Neafsey *et al.* 2015). To do this, I aligned the principal dipteran bait set to the genomes of additional dipteran lineages and an outgroup assembly from *Limnephilus lunatus* (GCA_000648945.1; Supplemental Table 3). Then, I sliced FASTA sequences from each genome that flanked the conserved locus location by ± 400 bp and performed additional orthology and duplicate screening steps (--min_coverage 67, --min_identity 80). I then created one data set containing members of Culicomorpha with *Limnephilus lunatus* as an outgroup taxon, and I created a second data set containing members of the Drosophilidae with *Musca domestica* and *Lucilia cuprina* as outgroup taxa. For each data set, I aligned the FASTA slices for each locus across all taxa using *mafft*, trimmed resulting alignments using *gblocks*, created two dataset in which all alignments contained data for at least 70% of the taxa, and concatenated each data set into a PHYLIP supermatrices, which I analyzed using best ML (GTRGAMMA) and bootstrap searches in *RAxML* v8.0.19 (Stamatakis 2014). I reconciled the best ML tree with the bootstrap replicates using *RAxML* v8.0.19.

### Hemiptera

The probe design process for hemipterans (Supplemental File 4) was similar those previously described: I downloaded available genomes for hemipteran lineages (Supplemental Table 4), and I used *art* (Huang *et al.* 2012) to simulate paired-end, 100 bp reads from each genome at 2x coverage for conserved locus identification. I merged read pairs into a single file and aligned the merged reads to the genome sequence of *Diaphorina citri* (GCA_000475195.1; diaPsy1 hereafter) using *stampy* (Lunter & Goodson 2011) with a substitution rate of 0.05. After removing unaligned reads, I followed the workflow for dipterans. I queried the resulting table of alignment intervals and output a list of intervals shared by the base taxon and three of the five exemplar taxa. I designed a temporary bait set targeting these loci from the diaPsy1 genome, and re-aligned the temporary bait set to the available genomes of exemplar taxa representing hemipteran diversity (Supplemental Table 4), designed a set of hybrid bait sequences targeting each conserved locus from the genomes of each exemplar taxon, and filtered duplicate baits/loci from this set. I called the resulting file the principal hemipteran bait set.

I followed the same procedures described above to perform an *in silico* check of the bait set. The only differences were that I aligned the baits to the genomes of taxa I used to design the principal hemipteran bait set, as well as two additional genome-enabled hemipteran lineages and an outgroup thysanopteran genome, *Frankliniella occidentalis* (Supplemental Table 4). I also performed the additional orthology and duplicate screening steps with slightly stricter parameters (--min_coverage 80, --min_identity 80). After alignment and trimming using *gblocks*, I created a data set in which all alignments contained at least 7 of the 10 taxa (70% complete matrix), I concatenated these loci, and I analyzed the concatenated alignment using *RAxML* as described above.

### Lepidoptera

Similar to the groups above, I downloaded available genomes for five lepidopteran lineages (Supplemental Table 5), as well as the genome assembly for *Limnephilus lunatus*, a caddisfly (Order Trichoptera). Then, I simulated paired-end, 100 bp reads using *art* (Huang *et al.* 2012). I merged read pairs into a single file for alignment to the *Bombyx mori* genome (GCA_000151625.1; bomMor1 hereafter) using *stampy* (Lunter & Goodson 2011) with a substitution rate of 0.05. After removing unaligned reads, I followed the workflow for dipterans, except that I did not merge aligned reads that were < 100 bp from one another. I queried the table of alignment intervals and output a list of intervals shared by the base taxon and all five of the exemplar taxa. I designed a temporary bait set from bomMor1 to target these loci, and I aligned the temporary bait set to the available genomes of exemplar taxa representing lepidopteran diversity (Supplemental Table 5). From these matches, I designed a set of hybrid bait sequences targeting each conserved locus from the genomes of each exemplar taxon, and I filtered duplicate baits/loci from this set. I called the resulting file the principal lepidopteran bait set.

I performed an *in silico* check of the principal lepidopteran bait set following the same procedures described above. In addition to re-aligning baits to the genomes of taxa I used for the bait set design, I included genome assemblies from 15 lepidopterans as well as the outgroup assembly from *Limnephilus lunatus* (Supplemental Table 5). I performed the orthology and duplicated screening steps with parameters identical to those used for dipterans, and after alignment and alignment trimming using gblocks, I created a data set in which all alignments contained 12 of the 16 taxa (75% complete matrix). I concatenated those loci and analyzed the concatenated matrix using RAxML, as described above.

## Results

### Arachnida

I collected an average of 39 M (95 CI: 7.3 M) sequencing reads from each low-coverage arachnid libraries (Supplemental Table 1). An average of 1.45% (95 CI: 0.8%) reads aligned to the limPol1 base genome sequence. After converting the alignments to BED format, merging overlapping alignment regions, and filtering BEDs of short loci or loci that aligned to large repeat regions in the limPol1 genome assembly, I selected 5,975 loci from the relational database that were shared by limPol1 and all six arachnid exemplars used for conserved locus identification. I designed a temporary bait set targeting 5,733 loci of these loci identified in the limPol1 genome assembly, and I re-aligned the temporary baits to the genomes of nine arachnids and limPol1. I selected a set of 1,168 conserved loci that were shared by limPol1 and at least five of the nine exemplar arachnid taxa, and I designed a hybrid bait set targeting these loci using the genomes of all nine arachnids and limPol1. After bait design and duplicate filtering, the principal arachnid probe set contained 14,799 baits targeting 1,120 loci.

During *in-silico* testing, I detected an average of 1,029 conserved loci among arachnid genome assemblies and the outgroup (limPol1) genome assembly, while the average number of non-duplicate, conserved loci was 692.8 (95 CI: 59.1). The 70% complete matrix contained 550 trimmed alignments that were 399 bp in length (95 CI: 16.73), totaled 219,372 characters, and contained 99,882 informative sites (mean per locus ± 95 CI: 182 ± 8). The resulting ML phylogeny (Supplemental Figure 1) reconstructed the established orders as monophyletic while recovering recognized relationships within spiders (Garrison *et al.* 2016) with high support at all nodes. Additional details regarding *in vitro* tests of this bait set can be found in (Starrett *et al.* Submitted).

### Coleoptera

I simulated an average of 7.0 M (95 CI: 3.8 M) sequencing reads from each coleopteran genome assembly (Supplemental Table 2), and approximately 1.2% (95CI: 0.2%) of these reads aligned to the triCas1 genome. After converting the alignments to BED format, merging overlapping regions, and filtering BEDs of short loci or loci that overlapped repetitive regions in the triCas1 genome, I selected 1,822 loci from the relational database that were shared by triCas1 and five exemplar taxa. I designed a temporary bait set from the triCas1 genome targeting 1,805 conserved loci, and I aligned the temporary baits to the genomes of six coleopterans and the strepsipteran outgroup. I selected a set of 1,209 conserved loci that were shared by triCas1 and at least four of the coleopteran and strepsipteran exemplar taxa, and I designed a hybrid bait set targeting these loci using the genomes of all seven coleopteran lineages and one strepsipteran lineage. The principal coleopteran bait set contained 13,674 baits targeting 1,172 conserved loci.

During *in-silico* testing, I detected an average of 994 conserved loci among coleopteran genome assemblies and the strepsipteran outgroup assembly, while the average number of non-duplicate, conserved loci detected in each taxon was 837.7 (95 CI: 105.9). After alignment and alignment trimming, the 70% complete matrix contained 865 loci that were 626.9 (95 CI: 9.8) bp in length, totaled 542,324 characters, and contained 163,681 informative sites (mean per locus ± 95 CI: 189.2 ± 3.6). The resulting ML phylogeny reconstructed recognized relationships among coleopteran superfamilies (Mckenna *et al.* 2015) with high support at all nodes (Supplemental Figure 2).

### Diptera

I simulated 2.9 M (95 CI: 0.3 M) sequencing reads from each of two dipteran genome assemblies (Supplemental Table 3). An average of 2.1% (95CI: 1.8%) of these reads aligned to the aedAeg1 genome. After converting the alignments to BED format, merging overlapping reads, and filtering BEDs of short loci or loci that overlapped repetitive regions in the aedAeg1 genome, I selected 4,904 conserved loci from the relational database that were shared by aedAeg1 and the two exemplar taxa. I designed a temporary bait set targeting these loci using the aedAeg1 genome assembly, and I aligned the temporary baits to the genomes of seven dipterans. I selected a set of 2,834 conserved loci that were shared by aedAeg1 and at least four other dipteran genome assemblies, and I designed a hybrid bait set targeting these loci using the genomes of seven dipteran lineages. The principal dipteran bait set contained 31,328 baits targeting 2,711 conserved loci.

During *in-silico* testing, I detected an average of 2,413 conserved loci among dipteran genome assemblies and the trichopteran outgroup assembly, while the average number of non-duplicate, conserved loci detected in each taxon was 1774.0 (95 CI: 213.6). I constructed two phylogenetic data matrices and inferred phylogenies from each. The 75% complete matrix for Culicomorpha contained 1,202 loci that were 676.4 (95 CI: 11.2) bp in length, totaled 813,084 characters, and contained 266,806 informative sites (mean per locus ± 95 CI: 222.0 ± 4.0). The resulting ML phylogeny (Supplemental Figure 3a) reconstructed the relationships among major mosquito/black fly lineages with high support at all nodes (Wiegmann *et al.* 2011), and the best ML topology was identical to a tree inferred from whole-genome sequence data (Neafsey *et al.* 2015). The 75% complete matrix for Drosophilidae contained 1,658 loci that were 721.2 (95 CI: 8.3) bp in length, totalled 1,195,791 characters, and contained 471,185 informative sites (mean per locus ± 95 CI: 284.2 ± 3.5). The resulting ML phylogeny (Supplemental Figure 3b) reconstructed the relationships among and within drosophilid lineages with high support at all nodes, and the best ML topology was similar to those of other studies (van der Linde *et al.* 2010; Wiegmann *et al.* 2011; Neafsey *et al.* 2015). The primary difference in topology between the conserved element tree and topologies inferred by other studies was the placement of *D. willistoni* sister to the *virilis+repleta+grimshawi* groups + subgenus Sophophora, a difference that could be explained by rooting the conserved element tree on *M. domestica*+*L. cuprina*.

### Hemiptera

I simulated 14.2 M (95 CI: 4.7 M) sequencing reads from each of the hemipteran genome assemblies (Supplemental Table 4). An average of 0.7% (95 CI: 0.3%) of these reads aligned to the diaPsy1 genome assembly. After converting the loci to BED format, merging overlapping reads, and removing short and repetitive loci, I selected 6,210 loci from the relational database that were shared by diaPsy1 and three exemplar taxa. I designed a temporary bait set targeting these loci from the diaPsy1 genome assembly, and I aligned the temporary baits to the genomes of eight hemipterans. I selected a set of 2,878 conserved loci shared by diaPsy1 and at least five of the hemipteran genome assemblies, and I designed a hybrid bait set targeting these loci using the genome assemblies of nine hemipteran lineages. The principal hemipteran bait set contained 40,207 baits targeting 2,731 conserved loci.

During *in-silico* testing, I detected an average of 2,381 conserved loci among hemipteran genome assemblies and the thysanopteran outgroup assembly, while the average number of non-duplicate, conserved loci detected in each taxon was 1,673.8 (95 CI: 223.1). The 75% complete matrix contained 1,444 loci that were 386.4 (95 CI: 7.1) bp in length, and the concatenated data matrix contained 557,988 characters and 260,127 informative sites (mean per locus ± 95 CI: 180.1 ± 3.1). The resulting ML phylogeny (Supplemental Figure 4) reconstructed recognized relationships among hemipteran lineages (Cryan & Urban 2012), particularly those within Heteroptera (Wang *et al.* 2016), with high support.

### Lepidoptera

I simulated an average of 6.9M (95 CI: 1.3 M) sequencing reads from each of the lepidopteran genome assemblies (Supplemental Table 5). An average of 4% (95CI: 1.2%) of these reads aligned to the bomMor1 base genome sequence. After converting the alignments to BED format, merging overlapping alignment regions, and filtering BEDs of short loci or loci that aligned to large repeat regions in the bomMor1 genome, I selected 2,162 conserved loci from the relational database that were shared among bomMor1 and the five exemplar taxa used for conserved region identification. I designed a temporary bait set containing 4,181 baits targeting 2,120 loci in bomMor1, and aligned that to the genome sequence of each exemplar taxon. I selected a set of 1,417 conserved loci that were shared by bomMor1 and at least three of the five exemplar taxa, and I designed a hybrid bait set targeting these loci using the genome assemblies of six lepidopteran lineages. After designing the baits and filtering duplicates, the principal lepidopteran bait set contained 14,363 baits targeting 1,381 conserved loci.

During *in-silico* testing, I detected an average of 1,141 conserved loci among lepidopteran genome assemblies and the trichopteran outgroup assembly, while the average number of non-duplicate, conserved loci detected in each taxon was 920.6 (95 CI: 39.9). The 75% complete matrix contained 876 conserved loci that were 463.3 (95 CI: 17.2) bp in length, and the concatenated data matrix contained 405,849 characters and 158,187 informative sites (mean per locus ± 95 CI: 180.5 ± 7.1). The resulting ML phylogeny (Supplemental Figure 5) reconstructed lepidopteran relationships that largely agree with recent phylogenomic studies (Kawahara and Breinholt 2014; Cong et al. 2015). Relationships within Papilionoidea do not differ from other studies. However, the placement of Pyraloidea sister to Papilionoidea in the conserved element phylogeny conflicts with previous studies that suggest Pyraloidea is sister to Macroheterocera+Mimallonidae (Bazinet *et al.* 2013; Kawahara & Breinholt 2014). Bootstrap support for this relationship is low, and a post-hoc DensiTree (Bouckaert 2010) analysis of the bootstrap replicates (Supplemental Figure 6) suggests that one cause of the low support for this relationship, as well as a source of the low support for the node uniting the Macroheterocera, is instability regarding to the placement of the Pyraloidea in the concatenated phylogenetic analysis.

## Discussion

I created a generalized workflow for (1) identifying conserved sequences shared among divergent genomes and (2) designing enrichment baits to collect these conserved regions from DNA libraries for downstream phylogenetic and phylogeographic analyses. Application of this workflow to several diverse groups of arthropods suggests that the method identifies thousands of conserved loci shared among divergent taxa using a handful of relatively simple steps. *In silico* testing suggests that these enrichment baits can be used to collect data from hundreds of loci across entire organismal groups, and *in silico* results also suggest that each bait set can be extended to divergent outgroups with moderate success. *In vitro* testing of the bait set designed for arachnids (Starrett *et al.* Submitted) suggests that *in silico* tests provide a reasonably accurate measure of success when baits are used to collect sequence data from real DNA libraries.

However, as with all newly designed target enrichment bait sets, readers should be cautioned that every bait set is “experimental” until validated *in vitro*. The number of bait sets I designed using the workflow described above puts *in vitro* testing of all of the arthropod bait sets beyond the scope of this manuscript, but each bait set is available, restriction-free under a public domain license (CC-0). This means that individual labs interested in testing remaining bait sets are free to use, modify, and extend any of the arthropod bait sets I have developed.

The workflow presented here differs from related efforts by combining target locus identification with enrichment bait design (Mayer *et al.* 2016) and because the process of conserved locus identification that I used is agnostic to the class of loci being interrogated (Johnson *et al.* 2016). This means that the conserved loci identified by the workflow I describe can be exons, introns, or intergenic regions. The set of conserved loci can be further subdivided into different classes using annotation information available from genomes to which the bait set is aligned or other data, such as transcript sequences. Furthermore, different algorithms for bait sequence selection and bait design (Mayer *et al.* 2016) can be applied to the conserved regions identified by the workflow I created to find improved or optimal bait designs.

Finally, because the workflow is generalized, its application is not limited to specific vertebrate or invertebrate classes - any organismal group having some genomic resources can be used for locus identification and subsequent bait design. And, the locus identification process can be tailored by users to be more or less strict than the moderate approach I used for each arthropod group, a strategy that allows researchers to identify variable numbers of conserved loci shared among focal taxa that scales with the risk each research group is willing to accept. For example, targeting those few hundred loci found in six out of six divergent taxa representing a given organismal group is less risky than targeting those few thousand loci that are putatively shared by only three of six divergent taxa.

By making all of the design steps, documentation, software code, and probe sets developed here available under an open-source license, I hope that the workflow I described will facilitate the collection of genome scale data from a diversity of organismal groups and provide additional insight into common and different patterns of diversification we see across the Tree of Life.

## Acknowledgements

I would like to thank the genome sequencing centers and individual research groups who make their genome assemblies publicly available - the work I described above would not be possible without these phenomenally useful resources. I would like to specifically thank Richard K. Wilson and The Genome Institute, Washington University School of Medicine for providing access to the *Limulus polyphemus* genome assembly through NCBI Genbank. I would also like to thank the Baylor College of Medicine Human Genome Sequencing Center (http://www.hgsc.bcm.tmc.edu), who have made all of the data generated as part of the i5k Initiative (i5K Consortium 2013) available to others. G. Dasch provided early access to the *Amblyomma americanum* genome assembly. I thank A. Chase for his assistance with preparing DNA libraries for sequencing and M. Branstetter, R. Bryson, S. Derkarabetian, T. Glenn, M. Hedin, J. McCormack, N. Pourmand, and J. Starrett who have contributed, in various ways, to the development process described above. C. Carlton, M. Forthman, C. Mitter, K. Noble, and C. Weirauch provided comments on phylogenetic trees inferred during *in silico* tests for different organismal groups, although any errors in describing those trees are my own. This work was supported by startup funds from Louisiana State University, with additional computational support from NSF DEB-1242260. Parts of this work were also encouraged by DEB-1352978 to David O’Brochta, whose invitation to speak at the IGTRCN symposium in 2014 spurred me to design probes targeting conserved loci in different insect groups. This study was also supported, in part, by resources and technical expertise from the Georgia Advanced Computing Resource Center, a partnership between the University of Georgia’s Office of the Vice President for Research and Office of the Vice President for Information Technology. Other portions of this research were conducted with high performance computing resources provided by Louisiana State University (http://www.hpc.lsu.edu).

## Data Accessibility

The workflow described above uses programs released as part of the open source PHYLUCE package (v1.6+; https://www.github.com/faircloth-lab/phyluce/) as well as a number of 3rd-party software programs that are cited in the main text. A generalized tutorial implementing the methods described as part of this manuscript is available from http://phyluce.readthedocs.io/en/latest/tutorial-four.html.

Raw reads used to identify conserved loci in arachnids are available from NCBI PRJNA324685 or directory from the SRA (Supplemental Table 1). Supplemental data, including files describing the probe design process, the final probe sets designed for this manuscript, and data from *in silico* testing are available from Dryad http://dx.doi.org/10.5061/dryad.v0k4h. All principal probe sets are also available from FigShare http://dx.doi.org/10.6084/m9.figshare.c.3472383, where I will maintain updated/improved versions.

---

**Supplemental Table 1.** Arachnid species used for conserved locus identification, bait design, and *in silico* testing of the resulting bait design.

| Species | Short Name | Accession (NCBI) | Used for<br>Cons.<br>Locus ID | Used For<br>Probe<br>Design | Used for<br>In-silico<br>Analyses | Reads | % Reads<br>Aligned to<br>limPol1 | Cons. Loci<br>Recovered<br>In-Silico |
| --- | --- | --- | --- | --- | --- | --- | --- | --- |
| *Trithyreus pentapalpis* | hubPen | SRR3879972 | ✓ |  |  | 35,897,068 | 0.98 |  |
| *Atypoides riversi* | antRiv | SRR3879973 | ✓ |  |  | 37,110,190 | 0.69 |  |
| *Phrynus marginemaculatus* | phrMar | SRR3879964 | ✓ |  |  | 34,708,762 | 2.19 |  |
| *Cryptocellus goodnighti* | cryGoo | SRR3879970 | ✓ |  |  | 28,457,032 | 0.75 |  |
| *Mitopus morio* | mitMor | SRR3879969 | ✓ |  |  | 64,176,903 | 4.1 |  |
| *Bothriurus keyserlingi* | botKey | SRR3879966 | ✓ |  |  | 29,844,175 | 0.61 |  |
| *Pseudouroctonus apacheanus* | pseApa | SRR3879967 | ✓ |  |  | 47,826,185 | 1.17 |  |
| *Hadogenes troglodytes* | hadTro | SRR3879965 | ✓ |  |  | 41,908,765 | 0.55 |  |
| *Vaejovis deboerae* | vaeDeb | SRR3879968 | ✓ |  |  | 32,025,105 | 2.01 |  |
| *Ixodes scapularis* | ixoSca1 | GCA_000208615.1 | ✓ | ✓ | ✓ | 26,367,884 | 0.23 | 755 |
| ***Limulus polyphemus*\* †** | limPol1 | GCA_000517525.1 | ✓ | ✓ | ✓ |  |  | 559 |
| *Acanthoscurria geniculata* | acaGen1 | GCA_000661875.1 |  | ✓ | ✓ |  |  | 696 |
| *Centruroides exilicauda* | cenExi1 | GCA_000671375.1 |  | ✓ | ✓ |  |  | 736 |
| *Latrodectus hesperus* | latHes1 | GCA_000697925.1 |  | ✓ | ✓ |  |  | 608 |
| *Loxosceles reclusa* | loxRec1 | GCA_001188405.1 |  | ✓ | ✓ |  |  | 593 |
| *Mesobuthus martensii* | mesmar1 | GCA_000484575.1 |  | ✓ | ✓ |  |  | 732 |
| *Parasteatoda tepidariorum* | parTep1 | GCA_000365465.1 |  | ✓ | ✓ |  |  | 784 |
| *Stegodyphus mimosarum* | steMim1 | GCA_000611955.2 |  | ✓ | ✓ |  |  | 851 |
| *Amblyomma americanum* | ambAme1 | Accession pending |  | ✓ | ✓ |  |  | 614 |

\* I used this taxon as the base genome, against which to align the other sequence data.
† I used this taxon as the outgroup in *in-silico* analyses

**Supplemental Table 2.** Coleopteran species used for conserved locus identification, bait design, and *in silico* testing of the resulting bait design.

| Species | Short Name | Accession (NCBI) | Used for<br>Cons. Locus<br>ID | Used For<br>Probe<br>Design | Used for<br>In-silico<br>Analyses | Reads | % Reads<br>Aligned to<br>triCas1 | Cons. Loci<br>Recovered In-Silico |
| --- | --- | --- | --- | --- | --- | --- | --- | --- |
| *Anoplophora glabripennis* | anoGla1 | GCA_000390285.1 | ✓ | ✓ | ✓ | 11,617,334 | 1.42 | 927 |
| *Agrilus planipennis* | agrPla1 | GCA_000699045.1 | ✓ | ✓ | ✓ | 4,687,488 | 0.97 | 903 |
| *Leptinotarsa decemlineata* | lepDec1 | GCA_000500325.1 | ✓ | ✓ | ✓ | 12,024,330 | 0.89 | 900 |
| *Onthophagus taurus* | ontTau1 | GCA_000648695.1 | ✓ | ✓ | ✓ | 3,022,362 | 1.48 | 854 |
| *Dendroctonus ponderosae* | denPon1 | GCA_000355655.1 | ✓ | ✓ | ✓ | 3,797,616 | 1.23 | 819 |
| ***Tribolium castaneum*\*** | triCas1 | GCA_000002335.2 | ✓ | ✓ | ✓ |  |  | 934 |
| ***Mengenilla moldrzyki* †** | menMol1 | GCA_000281935.1 |  | ✓ | ✓ |  |  | 527 |

\* I used this taxon as the base genome, against which to align the other sequence data.
† I used this taxon as the outgroup in *in-silico* analyses

**Supplemental Table 3.**
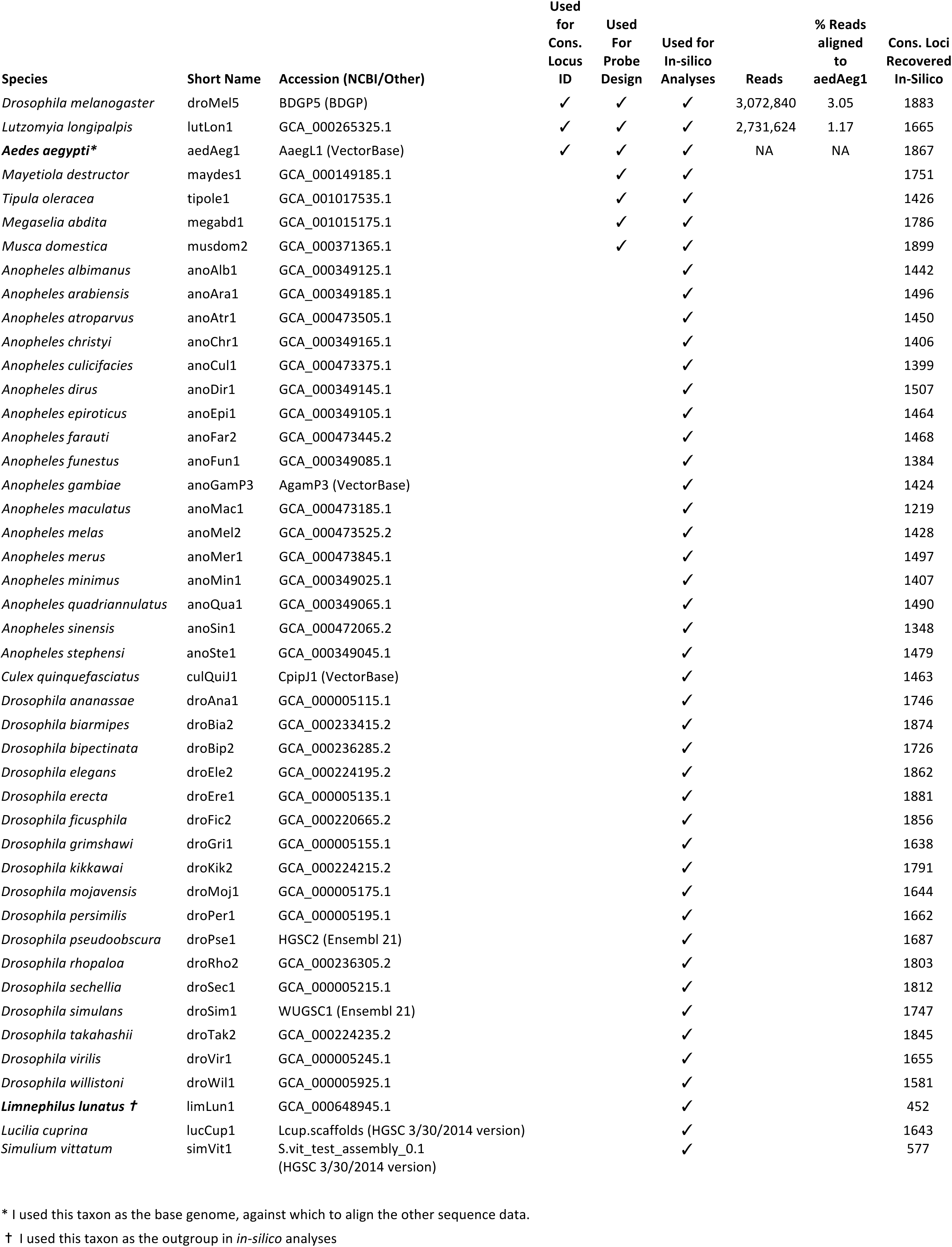
Dipteran species used for conserved locus identification, bait design, and in silico testing of the resulting bait design.

**Supplemental Table 4.** Hemipteran species used for conserved locus identification, bait design, and *in silico* testing of the resulting bait design.

| Species | Short Name | Accession (NCBI/Other) | Used for<br>Cons.<br>Locus ID | Used For<br>Probe<br>Design | Used for In-silico<br>Analyses | Reads | % Reads<br>aligned to<br>diaPsy1 | Cons. Loci<br>Recovered In-Silico |
| --- | --- | --- | --- | --- | --- | --- | --- | --- |
| *Acyrtosiphon pisum* | acyPis2 | GCA_000142985.2 | ✓ | ✓ | ✓ | 9,788,132 | 0.65 | 1414 |
| *Cimex lectularius* | cimLec1 | Clec_Bbug02212013 (HGSC) | ✓ | ✓ | ✓ | 10,051,932 | 1.48 | 1648 |
| *Gerris buenoi* | gerBue1 | GCA_001010745.1 | ✓ | ✓ | ✓ | 11,588,088 | 0.59 | 1908 |
| *Halyomorpha halys* | halHal1 | GCA_000696795.1 | ✓ | ✓ | ✓ | 19,393,098 | 0.43 | 1926 |
| *Homalodisca vitripennis* | homVit1 | GCA_000696855.1 | ✓ | ✓ | ✓ | 23,852,717 | 0.23 | 1914 |
| *Rhodnius prolixus* | rhoProC1 | RproC1 (VectorBase) | ✓ | ✓ | ✓ | 11,054,664 | 0.75 | 1919 |
| ***Diaphorina citri*\*** | diaPsy1 | GCA_000475195.1 | ✓ | ✓ | ✓ | NA | NA | 1369 |
| *Pachypsylla venusta* | pacVen1 | GCA_000695645.1 |  | ✓ | ✓ |  |  | 1786 |
| *Oncopeltus fasciatus* | oncFas1 | GCA_000696205.1 |  | ✓ | ✓ |  |  | 1990 |
| ***Frankliniella occidentalis* †** | fraOcc1 | GCA_000697945.1 |  |  | ✓ |  |  | 864 |

\* I used this taxon as the base genome, against which to align the other sequence data.
† I used this taxon as the outgroup in *in-silico* analyses

**Supplemental Table 5.** Lepidopteran species used for conserved locus identification, bait design, and *in silico* testing of the resulting bait design.

| Species | Short Name | Accession (NCBI) | Used for Cons. Locus ID | Used For Probe Design | Used for In-silico Analyses | Reads | % Reads aligned to bomMor1 | Cons. Loci Recovered In-Silico |
| --- | --- | --- | --- | --- | --- | --- | --- | --- |
| *Chilo suppressalis* | chiSup1 | GCA_000636095.1 | ✓ | ✓ | ✓ | 6,046,292 | 5.1 | 1074 |
| *Danaus plexippus* | danPle1 | GCA_000235995.1 | ✓ | ✓ | ✓ | 5,428,254 | 4.09 | 887 |
| *Heliconius melpomene* | helMel2 | GCA_000313835.2 | ✓ | ✓ | ✓ | 5,353,568 | 4.31 | 1042 |
| ***Limnephilus lunatus* †** | limLun1 | GCA_000648945.1 | ✓ | ✓ | ✓ | 13,572,210 | 2.18 | 936 |
| *Plutella xylostella* | pluXyl1 | GCA_000330985.1 | ✓ | ✓ | ✓ | 6,558,444 | - | 757 |
| ***Bombyx mori*\*** | bomMor1 | GCA_000151625.1 | ✓ | ✓ | ✓ | NA | NA | 1056 |
| *Amyelois transitella* | amyTra1 | GCA_001186105.1 |  |  | ✓ |  |  | 890 |
| *Lerema accius* | lerAcc1 | GCA_001278395.1 |  |  | ✓ |  |  | 923 |
| *Manduca sexta* | manSex1 | GCA_000262585.1 |  |  | ✓ |  |  | 946 |
| *Melitaea cinxia* | melCin1 | GCA_000716385.1 |  |  | ✓ |  |  | 882 |
| *Operophtera brumata* | opeBru1 | GCA_001266575.1 |  |  | ✓ |  |  | 836 |
| *Papilio glaucus* | papGla1 | GCA_000931545.1 |  |  | ✓ |  |  | 902 |
| *Papilio machaon* | papMac1 | GCA_001298355.1 |  |  | ✓ |  |  | 890 |
| *Papilio polytes* | papPol1 | GCA_000836215.1 |  |  | ✓ |  |  | 899 |
| *Papilio xuthus* | papXut1 | GCA_000836235.1 |  |  | ✓ |  |  | 938 |
| *Spodoptera frugiperda* | spoFru1 | GCA_000753635.2 |  |  | ✓ |  |  | 871 |

\* I used this taxon as the base genome, against which to align the other sequence data.
† I used this taxon as the outgroup in *in-silico* analyses

**Supplemental Figure 1.**
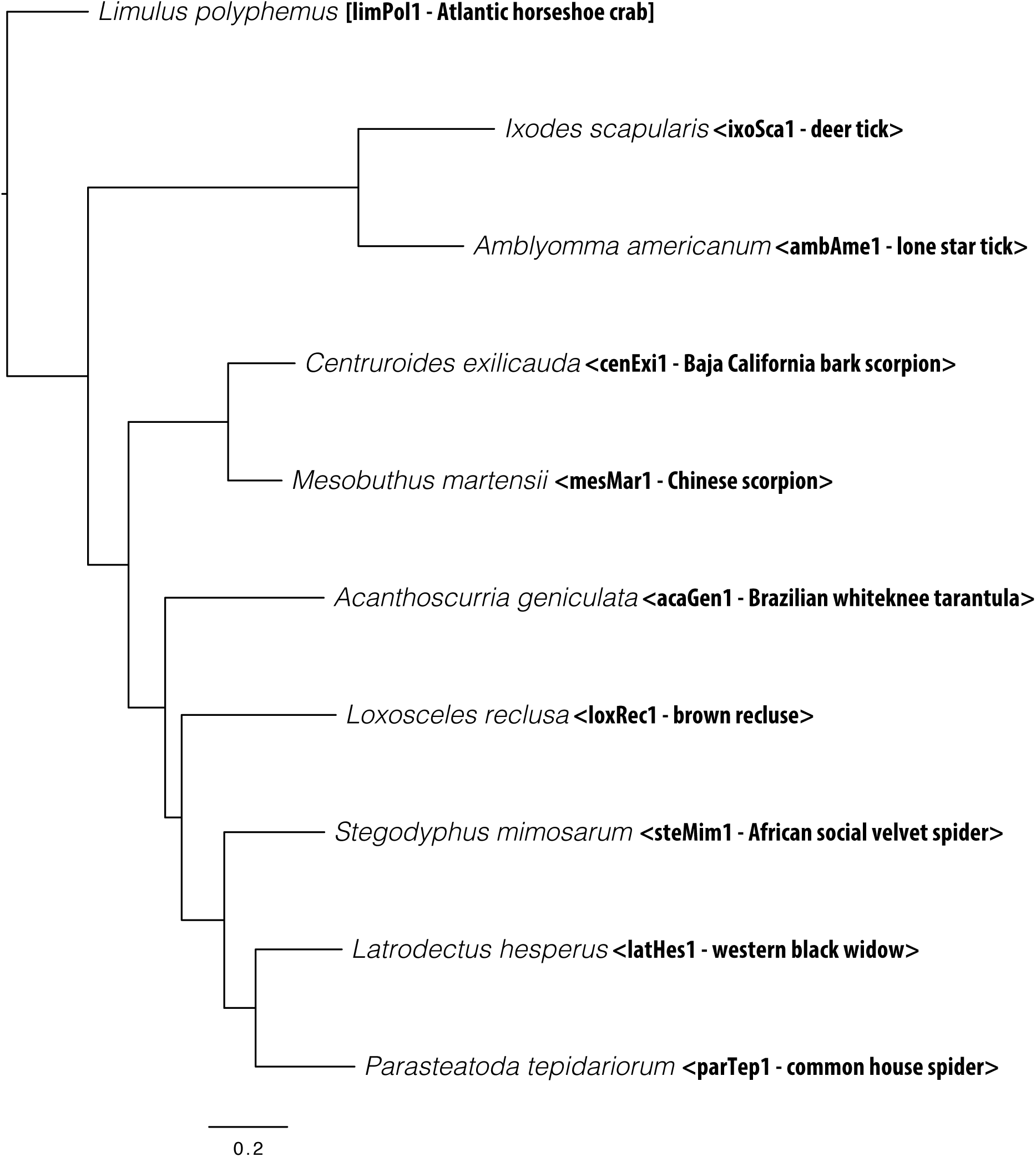
Maximum likelihood phylogeny inferred from *in silico* testing of baits targeting conserved loci in Arachnida. Bootstrap support values are 100% unless otherwise indicated.

**Supplemental Figure 2.**
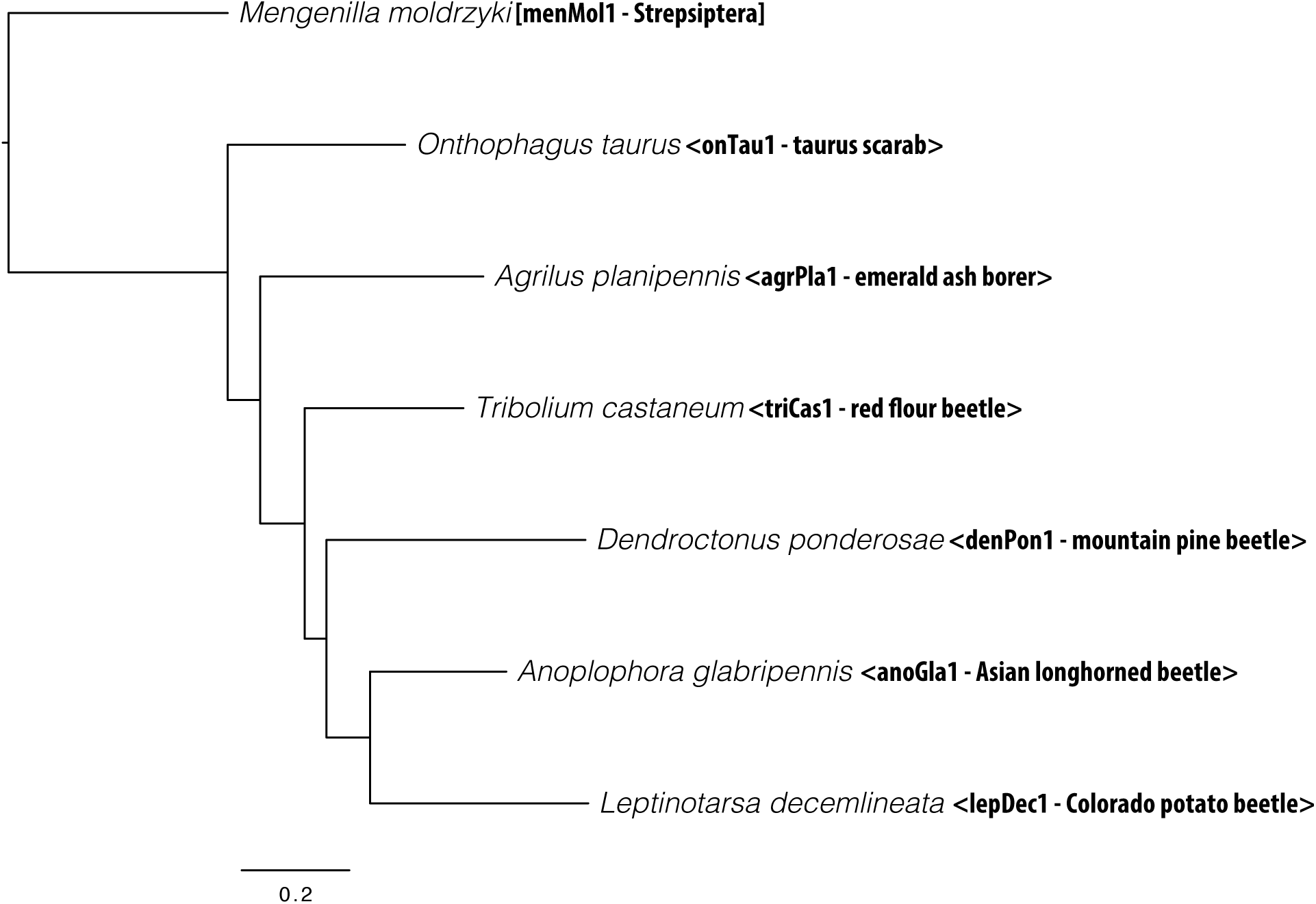
Maximum likelihood phylogeny inferred from *in silico* testing of baits targeting conserved loci in Coleoptera. Bootstrap support values are 100% unless otherwise indicated.

**Supplemental Figure 3.**
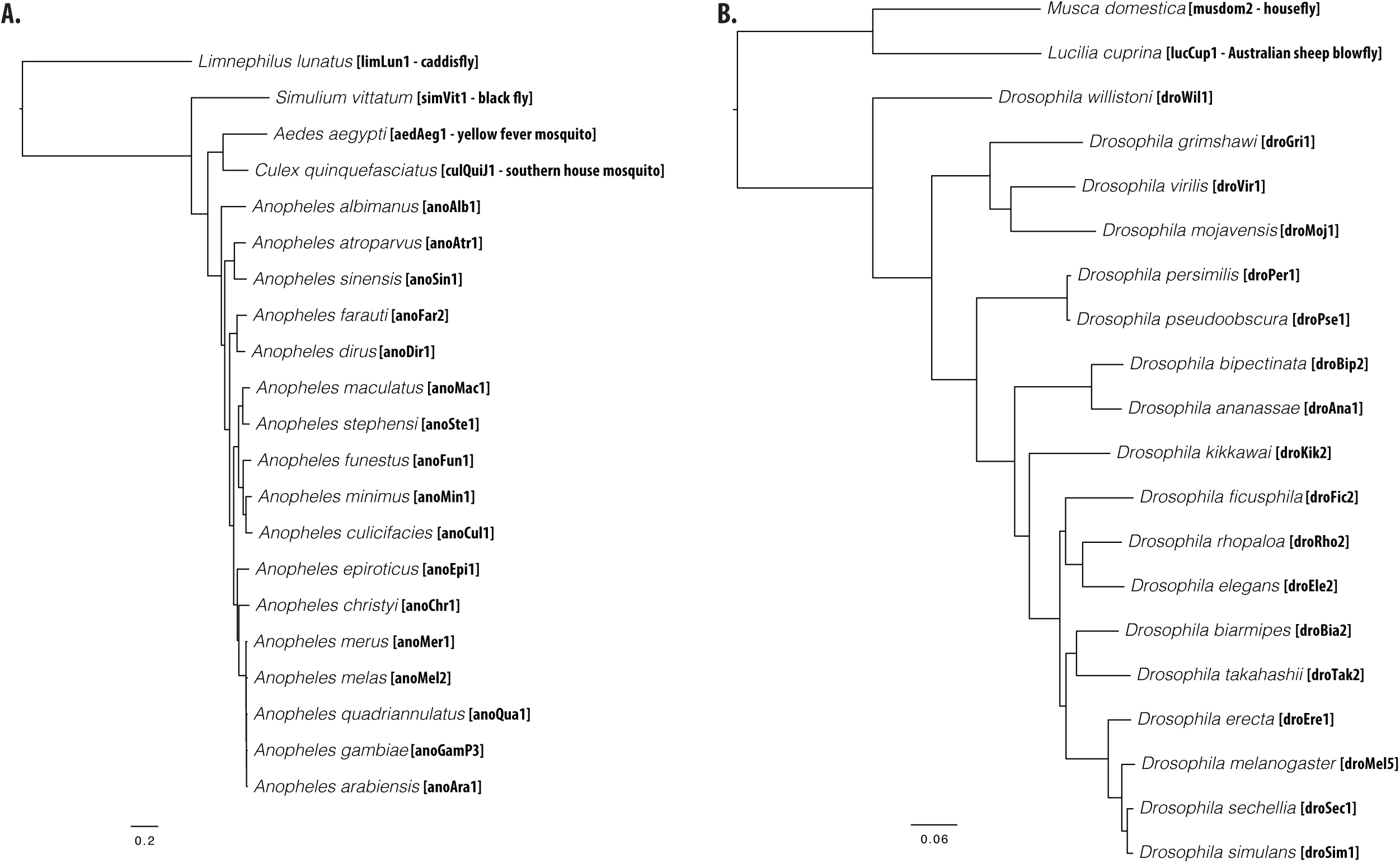
Maximum likelihood phylogeny inferred from *in silico* testing of baits targeting conserved loci in Diptera. Panel A shows relationships inferred among Culicomorpha and Panel B shoes relationships inferred among Drosophilidae. Bootstrap support values are 100% unless otherwise indicated.

**Supplemental Figure 4.**
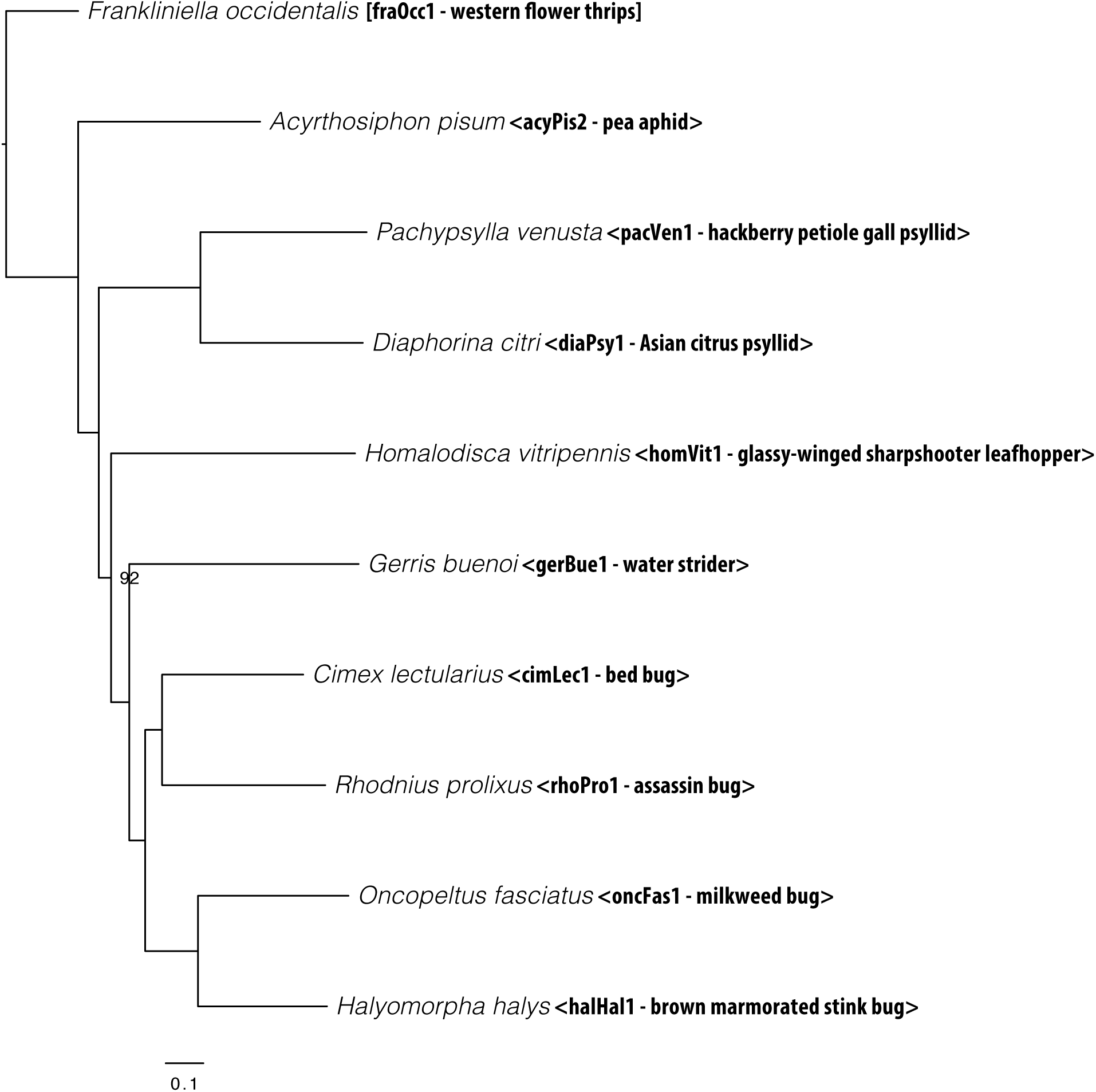
Maximum likelihood phylogeny inferred from in silico testing of baits targeting conserved loci in Hemiptera. Bootstrap support values are 100% unless otherwise indicated.

**Supplemental Figure 5.**
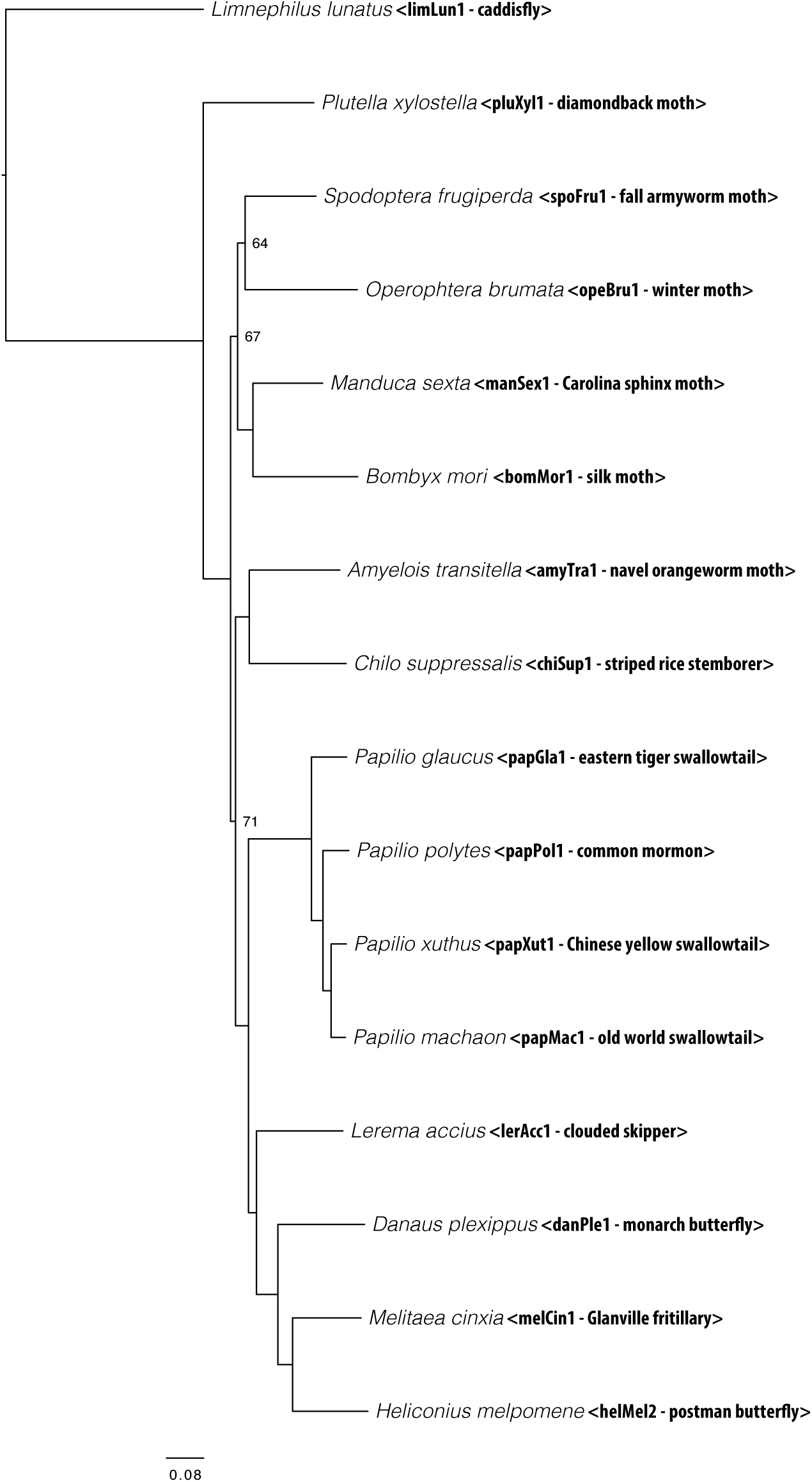
Maximum likelihood phylogeny inferred from in silico testing of baits targeting conserved loci in Lepidoptera. Bootstrap support values are 100% unless otherwise indicated.

**Supplemental Figure 6.**
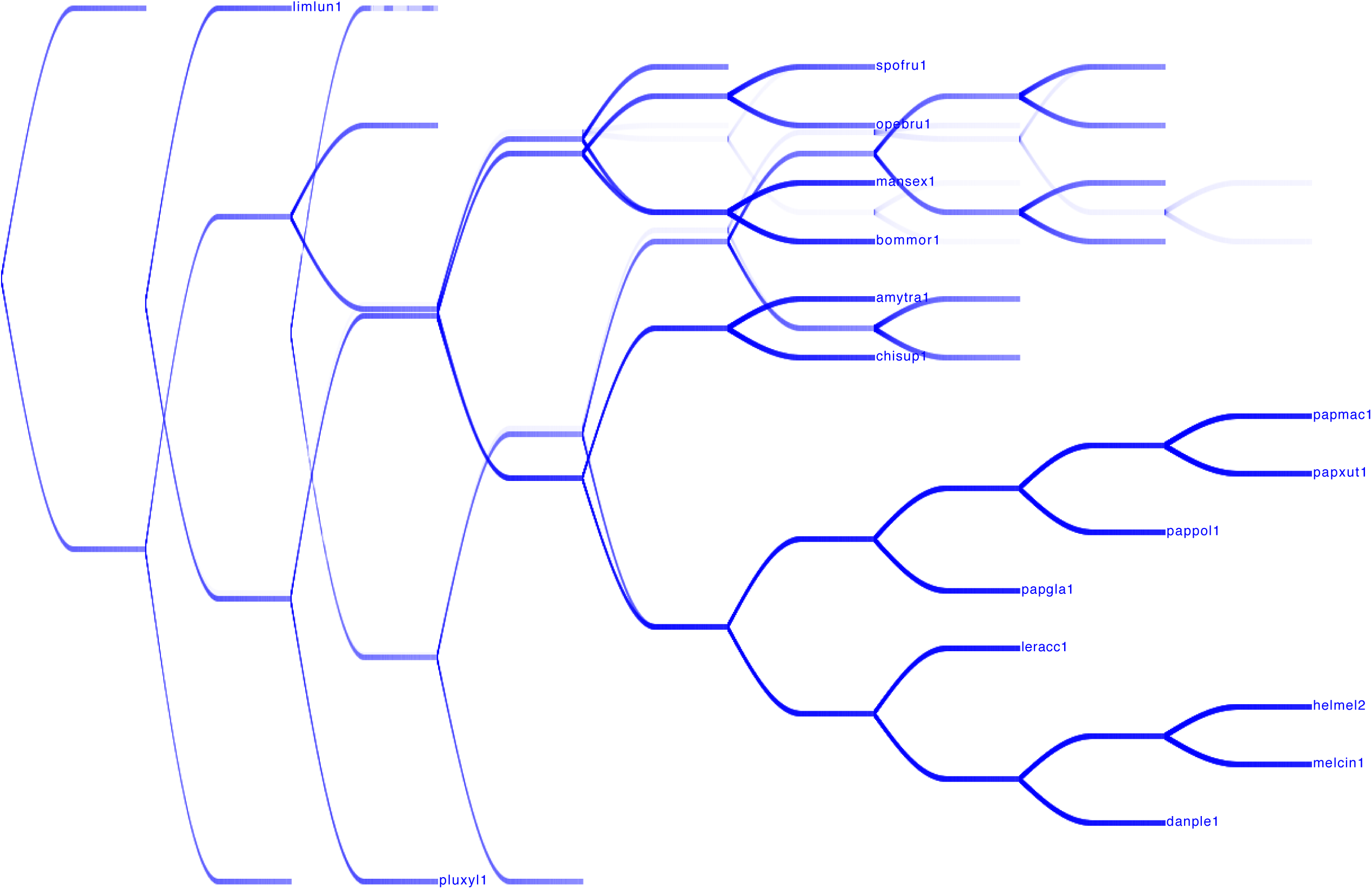
DensiTree plot of bootstrap replicates demonstrating instability regarding placement of the Pyraloidea in the concatenated phylogenetic analysis.

## References

Ali OA, O'Rourke SM, Amish SJ et al. (2015) RAD Capture (Rapture): Flexible and Efficient Sequence-Based Genotyping. Genetics, genetics.115.183665.

Baird N, Etter P, Atwood T et al. (2008) Rapid SNP discovery and genetic mapping using sequenced RAD markers. PloS one, 3, e3376.

Bazinet AL, Cummings MP, Mitter KT, Mitter CW (2013) Can RNA-Seq resolve the rapid radiation of advanced moths and butterflies (Hexapoda: Lepidoptera: Apoditrysia)? An exploratory study. PloS one, 8, e82615.

Bentley DR, Balasubramanian S, Swerdlow HP et al. (2008) Accurate whole human genome sequencing using reversible terminator chemistry. Nature, 456, 53–59.

Bi K, Linderoth T, Vanderpool D et al. (2013) Unlocking the vault: next-generation museum population genomics. Molecular ecology, 22, 6018–6032.

Bi K, Vanderpool D, Singhal S et al. (2012) Transcriptome-based exon capture enables highly cost-effective comparative genomic data collection at moderate evolutionary scales. BMC genomics, 13, 403.

Blaimer BB, Brady SG, Schultz TR et al. (2015) Phylogenomic methods outperform traditional multi-locus approaches in resolving deep evolutionary history: a case study of formicine ants. BMC evolutionary biology, 15, 271.

Blaimer BB, Lloyd MW, Guillory WX, Brady SG (2016) Sequence Capture and Phylogenetic Utility of Genomic Ultraconserved Elements Obtained from Pinned Insect Specimens. PloS one, 11, e0161531.

Bolger AM, Lohse M, Usadel B (2014) Trimmomatic: a flexible trimmer for Illumina sequence data. Bioinformatics, 30, 2114–2120.

Bouckaert RR (2010) DensiTree: making sense of sets of phylogenetic trees. Bioinformatics, 26, 1372–1373.

Bradnam KR, Fass JN, Alexandrov A et al. (2013) Assemblathon 2: evaluating de novo methods of genome assembly in three vertebrate species. GigaScience, 2, 10.

Castresana J (2000) Selection of conserved blocks from multiple alignments for their use in phylogenetic analysis. Molecular biology and evolution, 17, 540–552.

Crawford NG, Faircloth BC, McCormack JE et al. (2012) More than 1000 ultraconserved elements provide evidence that turtles are the sister group of archosaurs. Biology letters, 8, 783–786.

Cryan JR, Urban JM (2012) Higher!level phylogeny of the insect order Hemiptera: is Auchenorrhyncha really paraphyletic? Systematic entomology.

Drosophila 12 Genomes Consortium, Clark AG, Eisen MB et al. (2007) Evolution of genes and genomes on the Drosophila phylogeny. Nature, 450, 203–218.

Dunn CW, Hejnol A, Matus DQ et al. (2008) Broad phylogenomic sampling improves resolution of the animal tree of life. Nature, 452, 745–749.

Earl D, Bradnam K, St John J et al. (2011) Assemblathon 1: a competitive assessment of de novo short read assembly methods. Genome research, 21, 2224–2241.

Elshire RJ, Glaubitz JC, Sun Q et al. (2011) A robust, simple genotyping-by-sequencing (GBS) approach for high diversity species. PloS one, 6, e19379.

Faircloth BC (2013) illumiprocessor: a trimmomatic wrapper for parallel adapter and quality trimming.

Faircloth BC (2015) PHYLUCE is a software package for the analysis of conserved genomic loci. Bioinformatics, 32, 786–788.

Faircloth BC, Branstetter MG, White ND, Brady SG (2015) Target enrichment of ultraconserved elements from arthropods provides a genomic perspective on relationships among Hymenoptera. Molecular ecology resources, 15, 489–501.

Faircloth BC, Glenn TC (2012) Not all sequence tags are created equal: Designing and validating sequence identification tags robust to indels. PloS one, 7, e42543.

Faircloth BC, McCormack JE, Crawford NG et al. (2012) Ultraconserved elements anchor thousands of genetic markers spanning multiple evolutionary timescales. Systematic biology, 61, 717–726.

Faircloth BC, Sorenson L, Santini F, Alfaro ME (2013) A phylogenomic perspective on the radiation of ray-finned fishes based upon targeted sequencing of ultraconserved elements (UCEs). PloS one, 8, e65923.

Garrison NL, Rodriguez J, Agnarsson I et al. (2016) Spider phylogenomics: untangling the Spider Tree of Life. PeerJ, 4, e1719.

Gnirke A, Melnikov A, Maguire J et al. (2009) Solution hybrid selection with ultra-long oligonucleotides for massively parallel targeted sequencing. Nature biotechnology, 27, 182–189.

Hardenbol P, Banér J, Jain M et al. (2003) Multiplexed genotyping with sequence-tagged molecular inversion probes. Nature biotechnology, 21, 673–678.

Harris RS (2007) Improved pairwise alignment of genomic DNA. Ph.D. Thesis. The Pennsylvania State University.

Harvey MS (2002) The neglected cousins: what do we know about the smaller arachnid orders? The Journal of arachnology, 30, 357–372.

Harvey MG, Smith BT, Glenn TC, Faircloth BC, Brumfield RT (2016) Sequence capture versus restriction site associated DNA sequencing for shallow systematics. Systematic biology, 65, 910–924.

Hoffberg S, Kieran TJ, Catchen JM et al. (2016) RADcap: Sequence Capture of Dual-digest RADseq Libraries with Identifiable Duplicates and Reduced Missing Data. bioRxiv, 16, 1264–1278.

Hosner PA, Faircloth BC, Glenn TC, Braun EL, Kimball RT (2015) Avoiding missing data biases in phylogenomic inference: an empirical study in the landfowl (Aves: Galliformes). Molecular biology and evolution, 33, 1110–1125.

Huang W, Li L, Myers JR, Marth GT (2012) ART: a next-generation sequencing read simulator. Bioinformatics, 28, 593–594.

Hugall AF, O’Hara TD, Hunjan S, Nilsen R, Moussalli A (2016) An Exon-Capture System for the Entire Class Ophiuroidea. Molecular biology and evolution, 33, 281–294.

i5K Consortium (2013) The i5K Initiative: Advancing Arthropod Genomics for Knowledge, Human Health, Agriculture, and the Environment. The Journal of heredity, 104, 595–600.

Johnson MG, Gardner EM, Liu Y et al. (2016) HybPiper: Extracting coding sequence and introns for phylogenetics from high-throughput sequencing reads using target enrichment. Applications in plant sciences, 4.

Katoh K, Standley DM (2013) MAFFT multiple sequence alignment software version 7: improvements in performance and usability. Molecular biology and evolution, 30, 772–780.

Kawahara AY, Breinholt JW (2014) Phylogenomics provides strong evidence for relationships of butterflies and moths. Proceedings. Biological sciences /The Royal Society, 281, 20140970.

Li H, Handsaker B, Wysoker A et al. (2009) The Sequence Alignment/Map format and SAMtools. Bioinformatics, 25, 2078–2079.

Lim HC, Braun MJ (2016) High!throughput SNP genotyping of historical and modern samples of five bird species via sequence capture of ultraconserved elements. Molecular ecology resources, 16, 1204–1223.

van der Linde K, Houle D, Spicer GS, Steppan SJ (2010) A supermatrix-based molecular phylogeny of the family Drosophilidae. Genetics research, 92, 25–38.

Lunter G, Goodson M (2011) Stampy: a statistical algorithm for sensitive and fast mapping of Illumina sequence reads. Genome research, 21, 936–939.

Manthey JD, Campillo LC, Burns KJ, Moyle RG (2016) Comparison of Target-Capture and Restriction-Site Associated DNA Sequencing for Phylogenomics: A Test in Cardinalid Tanagers (Aves, Genus: Piranga). Systematic biology, 65, 640–650.

Mayer C, Sann M, Donath A et al. (2016) BaitFisher: A Software Package for Multispecies Target DNA Enrichment Probe Design. Molecular biology and evolution, 33, 1875–1886.

McCormack JE, Faircloth BC, Crawford NG et al. (2012) Ultraconserved elements are novel phylogenomic markers that resolve placental mammal phylogeny when combined with species tree analysis. Genome research, 22, 746–754.

McCormack JE, Harvey MG, Faircloth BC et al. (2013) A phylogeny of birds based on over 1,500 loci collected by target enrichment and high-throughput sequencing. PloS one, 8, e54848.

McCormack JE, Tsai WLE, Faircloth BC (2015) Sequence capture of ultraconserved elements from bird museum specimens. Molecular ecology resources, 16, 1189–1203.

McGee MD, Faircloth BC, Borstein SR et al. (2016) Replicated divergence in cichlid radiations mirrors a major vertebrate innovation. Proceedings. Biological sciences /The Royal Society, 283.

Mckenna DD, Wild AL, Kanda K (2015) The beetle tree of life reveals that Coleoptera survived end!Permian mass extinction to diversify during the Cretaceous terrestrial revolution. Systematic Entomology, 40, 835–880.

Misof B, Liu S, Meusemann K et al. (2014) Phylogenomics resolves the timing and pattern of insect evolution. Science, 346, 763–767.

Neafsey DE, Waterhouse RM, Abai MR et al. (2015) Mosquito genomics. Highly evolvable malaria vectors: the genomes of 16 Anopheles mosquitoes. Science, 347, 1258522.

Peñalba JV, Smith LL, Tonione MA et al. (2014) Sequence capture using PCR-generated probes: a cost-effective method of targeted high-throughput sequencing for nonmodel organisms. Molecular ecology resources, 14, 1000–1010.

Peterson BK, Weber JN, Kay EH, Fisher HS, Hoekstra HE (2012) Double digest RADseq: an inexpensive method for de novo snp discovery and genotyping in model and non-model species. PloS one, 7, e37135.

Quinlan AR, Hall IM (2010) BEDTools: a flexible suite of utilities for comparing genomic features. Bioinformatics, 26, 841–842.

Rohland N, Reich D (2012) Cost-effective, high-throughput DNA sequencing libraries for multiplexed target capture. Genome research, 22, 939–946.

Smith BT, Harvey MG, Faircloth BC, Glenn TC, Brumfield RT (2014) Target capture and massively parallel sequencing of ultraconserved elements (UCEs) for comparative studies at shallow evolutionary time scales. Systematic biology, 63, 83–95.

Smith SA, Wilson NG, Goetz FE et al. (2011) Resolving the evolutionary relationships of molluscs with phylogenomic tools. Nature, 480, 364–367.

Stamatakis A (2014) RAxML version 8: a tool for phylogenetic analysis and post-analysis of large phylogenies. Bioinformatics, 30, 1312–1313.

Starrett J, Derkarabetian S, Hedin M et al. (Submitted) High Phylogenetic Utility of an Ultraconserved Element Probe Set Designed for Arachnida. Molecular ecology resources.

Streicher JW, Wiens JJ (2016) Phylogenomic analyses reveal novel relationships among snake families. Molecular phylogenetics and evolution, 100, 160–169.

Suchan T, Pitteloud C, Gerasimova NS et al. (2016) Hybridization Capture Using RAD Probes (hyRAD), a New Tool for Performing Genomic Analyses on Collection Specimens. PloS one, 11, e0151651.

Talavera G, Castresana J (2007) Improvement of phylogenies after removing divergent and ambiguously aligned blocks from protein sequence alignments. Systematic biology, 56, 564–577.

Wang Y-H, Cui Y, Rédei D et al. (2016) Phylogenetic divergences of the true bugs (Insecta: Hemiptera: Heteroptera), with emphasis on the aquatic lineages: the last piece of the aquatic insect jigsaw originated in the Late Permian/Early Triassic. Cladistics: the international journal of the Willi Hennig Society, 32, 390–405.

Wiegmann BM, Trautwein MD, Winkler IS et al. (2011) Episodic radiations in the fly tree of life. Proceedings of the National Academy of Sciences of the United States of America, 108, 5690–5695.

Zhang Z-Q (2011) Phylum Arthropoda von Siebold, 1848. In: Zhang, Z.-Q.(Ed.) Animal. name Zootaxa, 3148, 99–103.

